# Bessel-droplet foci enable high-resolution and high-contrast volumetric imaging of synapses and circulation in the brain *in vivo*

**DOI:** 10.1101/2022.03.05.483143

**Authors:** Wei Chen, Qinrong Zhang, Ryan Natan, Jianglan Fan, Na Ji

## Abstract

Bessel beam has long been utilized in physics for its ability to maintain lateral confinement during propagation. When used for two-photon fluorescence microscopy, Bessel foci have enabled high-speed volumetric imaging of the brain. At high numeric aperture (NA), however, the substantial energy in the side rings of Bessel foci reduces image contrast. Therefore, a compromise between resolution and contrast has to be made, limiting Bessel foci in microscopy to low NA. Here, we describe a method of generating axially extended Bessel-droplet foci with much suppressed side rings. Shaping the excitation wavefront with novel phase patterns, we generated Bessel-droplet foci of variable NAs at high power throughput and scanned them interferometrically along the axial direction for continuous volume imaging. More resistant to optical aberrations than Bessel foci, Bessel-droplet foci enabled high-resolution and high-contrast volumetric imaging of synaptic anatomy and function as well as lymphatic circulation in the mouse brain *in vivo*.

## Introduction

Modern optical microscopy aims to probe biological samples at both high spatial and temporal resolutions volumetrically. This is of particular importance for imaging neurons and vessels in the brain, which are connected via synapses or capillaries into three dimensional (3D) functional systems with subsecond temporal dynamics (1–3). Among the major advances of optical microscopy in the past decades, two-photon fluorescence microscopy (2PFM) has revolutionized in vivo brain imaging by offering subcellular resolution in highly scattering brain tissues (4, 5), allowing the visualization of neuronal activity at synapses and cerebral circulation in capillaries in vivo (6, 7).

Traditional 2PFM scans a focus that is highly confined in 3D within the sample, and records the fluorescence generated at each position point-by-point to map the fluorophore distribution. To image neurons or vascular networks volumetrically, the excitation focus has to be scanned in 3D, which substantially reduces the achievable temporal resolution. Among the many high-speed volumetric imaging techniques developed for 2PFM (2, 8, 9), approaches that use a Bessel-like beam for two-photon excitation (10–15) stand out for its simplicity and ease in implementation in existing microscopy systems. Axially extended but laterally confined, when scanned in two dimensions (2D), a Bessel focus probes a 3D volume, generating a projected view of the volume at high temporal and spatial (in the lateral XY plane) resolution. Optimizing Bessel focus for in vivo imaging of neurons and vasculatures (13, 15–18), we have previously demonstrated high-speed volumetric two-photon imaging of synaptic and capillary activity across large volumes.

However, in addition to the central peak of the point spread function, the axially extending side rings of the Bessel focus can also excite fluorescence, which reduces image contrast. Particularly prominent in Bessel beams of high numeric aperture (NA), these side rings limited the excitation NA in previous studies to ≤0.4. Three-photon fluorescence microscopy has been combined with Bessel focus to suppress the contribution of the side rings, due to its higher order nonlinear excitation (19, 20). However, three-photon excitation requires expensive laser systems that are much less accessible to microscopy users.

Here, we describe a method of generating axially extended Bessel-droplet foci with substantially suppressed side rings by shaping the wavefront of the excitation light. Using novel phase patterns on a liquid-crystal spatial light modulator (SLM), we generated Bessel-droplet foci of variable NA and axial length at high power throughput and scanned them interferometrically along the axial direction for continuous volume imaging. When combined with 2PFM, Bessel-droplet foci led to a up to 10.6-fold reduction in side-ring fluorescence, enabling high-resolution imaging operations up to 0.7 NA. More resistant to optical aberrations than Bessel foci, Bessel-droplet foci enabled high-contrast, high-resolution volumetric imaging of synaptic anatomy and function as well as lymphatic circulation in the mouse brain in vivo. With the Bessel-droplet foci, we achieved volumetric 2PFM imaging via extended depth of focus at high NA with uncompromised contrast and resolution.

## Results

### Design of Bessel-droplet foci with minimal side rings for two-photon fluorescence microscopy

We constructed the Bessel-droplet module on a home-built 2PFM (21), where pairs of telecentric f-θ lenses (L4 – L7, Special Optics) optically conjugated two galvanometer mirrors (“galvos”) and the back pupil plane of the microscope objective (Olympus, 25×, 1.05 NA, 2 mm WD) (Fig. 1a). The 2PFM was designed to be switchable between two optical paths: one used the standard 2P excitation with a Gaussian illumination profile at the back pupil (yellow path, “Gaussian focus”); the other used an axially extended focus for 2P excitation (red path). For the latter, before reaching the galvos, the excitation light (940 nm, Insight DeepSee, Spectral Physics) reflected off a phase-only liquid crystal SLM (SLM, HSP1920, Meadowlark Optics) placed at a location that was conjugated to the objective focal plane. A lens (L1) Fourier-transformed the phase-modulated excitation light field onto an amplitude mask (chrome coating, Photo Sciences, Inc). Spatially filtering out stray and undiffracted light, the mask was conjugated to the galvos and objective pupil plane with two lenses (L2, L3), with the transmitted excitation electric field distribution designed to achieve the desired excitation point spread function (PSF). Another phase-only liquid crystal SLM (not shown) was located between and conjugated to the galvos and the objective (21), and can be used to shift the excitation focus axially by displaying a defocus pattern (22).

**Fig. 1.**
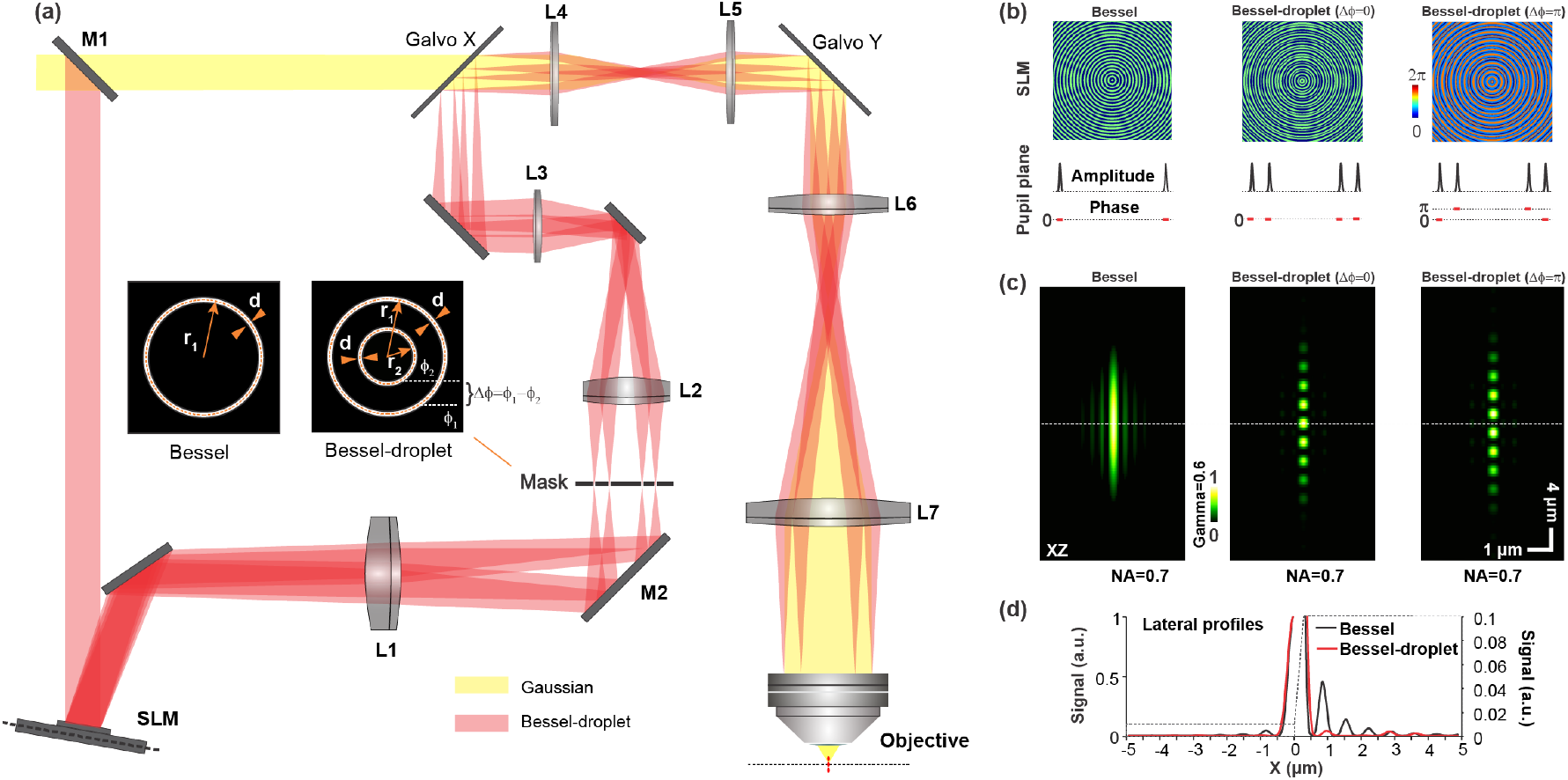
A 2PFM with a SLM-based module that generates axially extended Bessel-droplet foci with suppressed side rings. (**a**) Optical path diagram of the 2PFM. Gaussian path (yellow) and the path with a Bessel-droplet focus (red) are switchable via mirror M1. Insets: illumination patterns at the mask/objective back pupil plane for Bessel and Bessel-droplet foci, respectively. (**b**) (Top) Phase profiles on the SLM and (Bottom) the corresponding cross-sectional profiles of electric field amplitude and phase distributions at the mask/objective back pupil plane in order to generate (**c**) (From left to right, simulated axial PSFs of) a Bessel focus, a Bessel-droplet focus with 0 phase offset between two annuli (Δφ=0), a Bessel-droplet focus with π phase offset between two annuli (Δφ=π), all with NA 0.7. (d) Lateral profiles of the simulated Bessel and Bessel-droplet PSFs.

A conventional Bessel focus can be generated by an annular illumination at the objective back pupil plane (left inset, Fig. 1a). For an infinitely thin annulus, all rays post objective have the same axial components of wavevectors, which leads to the absence of axial intensity modulation and an infinitely long Bessel focus. Each ray within the annulus has a corresponding ray with the opposite lateral wavevector component; the interference between the two rays results in a standing wave in the lateral focal plane. All the standing waves from pairs of rays have the same spatial frequency and combine to form the characteristic lateral PSF of Bessel beam: a central peak surrounded by concentric side rings.

We can generate a Bessel-like focus by applying a concentric binary grating pattern of phase values 0 and π to the SLM (left panel, Fig. 1b), which diffracts most of the excitation light energy into an annulus at the mask/back pupil plane. We simulated a Bessel-like focus with 0.7 NA and 28 μm in axial full width at half maximum (FWHM) using a vectorial diffraction model (23). Even with suppression by two-photon excitation, its two-photon fluorescence PSF had substantial side rings (left panels, Fig. 1c; Fig. 1d). This is consistent with our earlier experimental measurements, where the more prominent side rings at higher NA caused substantial background excitation and reduced image contrast in the mouse brain (13). Consequently, Bessel focus used for 2PFM mostly had NA≤0.4 to limit side-ring contamination, which lowered lateral resolution and focal intensity.

Side rings can be reduced if two concentric annuli of illumination are used for excitation (right inset, Fig. 1a). The two annuli generate two sets of standing waves in the lateral focal plane with distinct spatial frequencies. When designed properly, they can interfere and suppress energy within the side rings. The distinct axial wave vectors from rays within the two annuli lead to modulation of axial intensity, with periodic low and high intensity regions in the axially extended PSF appearing as “droplets” of excitation. In other words, Bessel droplets can be considered as resulting from the interference of two Bessel beams, whose side rings can destructively interfere and reduce the energy distributed therein.

Because of the high-power requirement for two-photon excitation, we developed a computation procedure to calculate phase profiles that, when displayed by the SLM, converted the standard laser output with a Gaussian illumination profile into two concentric annuli at the mask/back pupil plane (middle and right panels, Fig. 1b; supplementary text 1 and fig. S1). For Bessel-droplet foci of NA 0.4-0.7, the optical throughput of our SLM-based module was above 35%, substantially higher than if the amplitude mask alone was used to transmit two annular regions of the laser illumination (supplementary text 2 and fig. S2).

A particularly novel aspect of our approach is that we could generate two illumination annuli of arbitrary phase offset Δϕ, which led to displacement of the Bessel droplets along the axial direction. In simulation, a phase profile designed to generate two annuli of π phase offset led to a complementary axial intensity profile (right panels, Fig. 1b,c) to that of the Bessel droplets formed by two annuli of the same phase (middle panels, Fig. 1b,c), with both Bessel-droplet foci of 0.7 NA exhibiting strong suppression of side rings in their simulated two-photon fluorescence PSF (Fig. 1c,d). Alternating the two phase profiles on the SLM during 2PFM imaging, we could rapidly scan the Bessel-droplet excitation axially. This provides a simple and elegant way to probe continuous volumes at high contrast and lateral resolution as detailed below.

The flexible phase control offered by the SLM enabled us to systematically explore the design space in order to find the concentric annular illumination patterns that maximally suppressed side rings of Bessel droplets. For both Bessel and Bessel-droplet foci, the thickness of the annuli *(d)* determines its axial extension. For Bessel focus, the radius of the annulus determines its NA; For Bessel-droplet focus, the radius of the outer annulus *(r1)* determines its NA. Holding *d* and *r_1_* (i.e., axial length and NA) constant, we systematically varied the inner annulus radius *(r_2_)* and simulated the two-photon excitation PSFs for NA 0.4, 0.5, 0.6, and 0.7 (23) (supplementary text 3, fig. S3). We then calculated the ratio between the peak of the most prominent side ring and the central peak of the lateral PSF, and found the optimal *r_2_* that minimized this ratio (fig. S3b). We further quantified the side-ring contamination by the ratio between the integrated signal within the most prominent side ring and the integrated signal within the central peak of the lateral PSF. At optimized *r_2_*’s (0.5*r_1_* – 0.6*r_1_*), we found this side-ring contamination ratio to be 0.036, 0.043, 0.048, and 0.048 for Bessel-droplet foci of NA 0.4, 0.5, 0.6, and 0.7, respectively (fig. S3d). In contrast, the side-ring contamination ratios for Bessel foci of the same NAs were 0.29, 0.31, 0.34, and 0.36, respectively (fig. S3c), 7 to 8 fold larger than Bessel-droplet foci. The strong suppression of the side rings throughout the range of NA can be easily appreciated by comparing the simulated axial and lateral PSFs of Bessel versus Bessel-droplet foci (fig. S3e).

### Experimental characterization of Bessel-droplet foci

From the concentric annular patterns that maximally suppressed side rings, we calculated and displayed the corresponding phase profiles on the SLM and measured the two-photon excitation PSFs of Bessel-droplet foci of NA 0.4, 0.5, 0.6, and 0.7 using 0.2-μm-diameter fluorescent beads. Comparing their axial and lateral PSFs with those of the Bessel foci of the same NA and annular thickness (Fig. 2a-c), we found that, consistent with the simulation results, Bessel foci had more prominent side rings at higher NA. In contrast, side rings of Bessel-droplet foci were greatly suppressed across the entire NA range, as clearly shown in the radially averaged profiles of the lateral PSFs (Fig. 2d). For Bessel-droplet foci of NA 0.4, 0.5, 0.6, and 0.7, the ratios between the integrated signal from the 1^st^ side ring and the integrated signal from the central peak of their lateral PSFs showed a 10.6-fold, 6.7-fold, 8.2-fold, 7.8-fold reduction, respectively, when compared to those of Bessel foci of the same NA. The lateral FWHMs of Bessel-droplet PSFs were measured at 0.74, 0.65, 0.60, and 0.55 μm for NA 0.4, 0.5, 0.6, and 0.7, respectively. In the spatial frequency domain, the resolution of Bessel-droplet foci were characterized from the cut-off frequencies of the optical transfer functions (calculated as the Fourier transform of PSFs) to be 0.70, 0.55, 0.47, and 0.42 μm for NA 0.4, 0.5, 0.6, and 0.7, respectively. The experimentally measured resolution was ~7% lower than those of Bessel foci of the same NA (0.68, 0.53, 0.44, and 0.37 μm for NA 0.4, 0.5, 0.6, and 0.7, respectively), consistent with theoretical calculation.

**Fig. 2.**
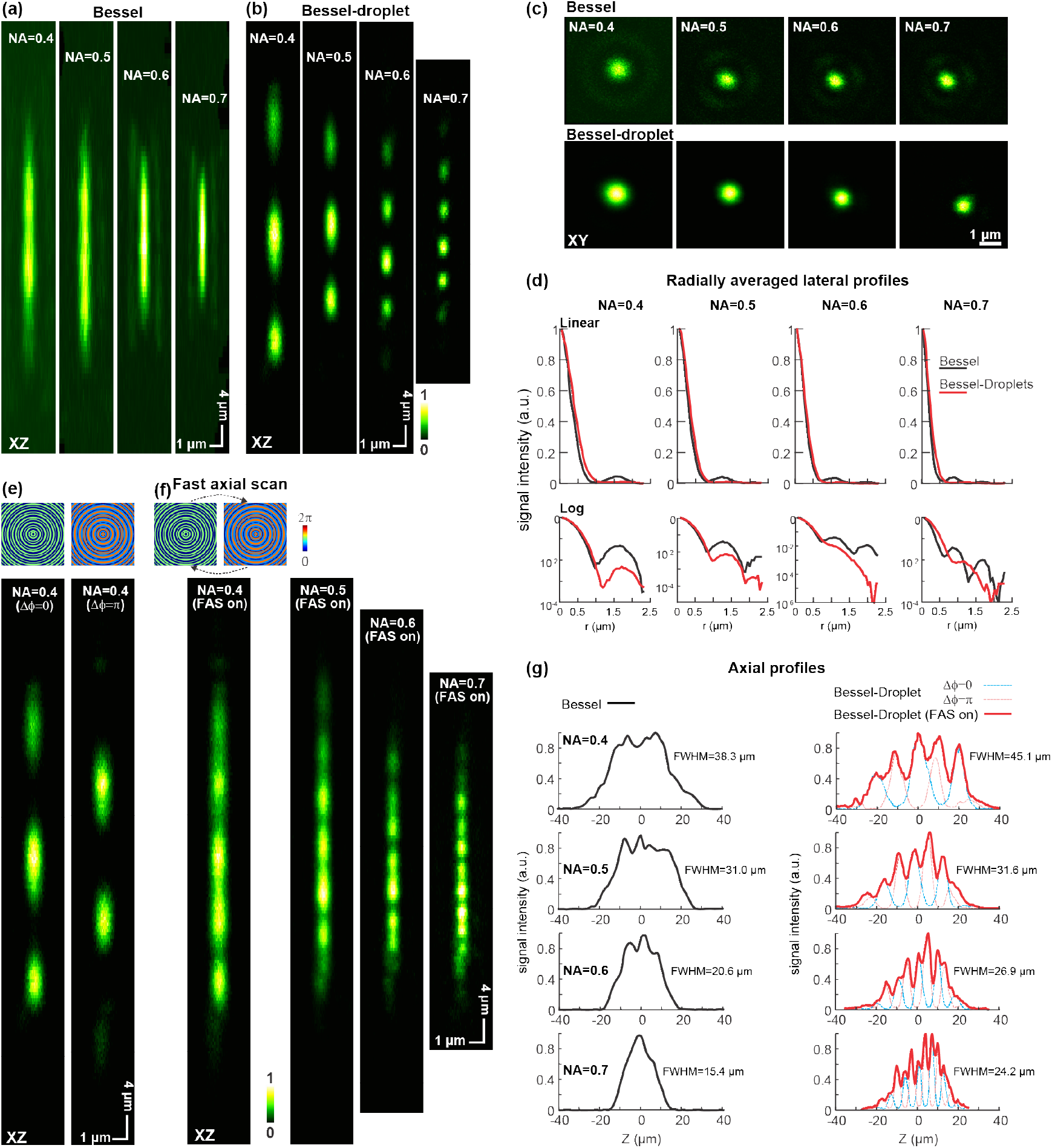
Experimentally measured 2PFM PSFs and fast axial scanning of Bessel and Bessel-droplet foci. (**a,b**) Axial PSFs of Bessel (**a**) and Bessel-droplet (**b**) foci of NA 0.4, 0.5, 0.6, 0.7. (**c**) Lateral PSFs of the foci in **a** and **b**. (**d**) Radially averaged profiles of the lateral PSFs in **c**, plotted in linear and semi-logarithmic scales. (**e**) Two phase patterns on SLM that give rise to 0 or π phase offset of the two annuli and the corresponding axial PSFs of Bessel-droplet foci of NA 0.4. (**f**) fast axial scanning (FAS) by switching between the two phase patterns enables continuous volume sampling, as shown by the composite axial PSFs of Bessel-droplet foci of NA 0.4, 0.5, 0.6, 0.7. (**g**) Axial profiles of Bessel PSFs in **a** and Bessel-droplet PSFs in **f.** Dashed lines: profiles of Bessel-droplet foci with Δϕ = 0, π. All PSFs measured using 0.2-μm-diameter beads with the same post-objective power and displayed in scales normalized to their own maximum signal, with optical system aberrations corrected following Ref. [24].

As predicted by simulation, by changing the phase profiles displayed on the SLM to generate two concentric annular illumination with the phase offset Δφ being 0 or π, we shifted the axial locations of the Bessel droplets so that they form complementary excitation profiles (Fig. 2e, supplementary text 4, fig. S4). Taking advantage of the high refreshing rate of our SLM (1.18 ms), we switched the SLM between the two phase profiles for every other image frame. With such a fast axial scanning (FAS) scheme, the entire depth extent of the Bessel-droplet foci can be probed without missing volumes (Fig. 2f). Alternatively, we applied defocus patterns on a second SLM conjugated to the objective back pupil plane to remotely scan the Bessel droplets axially between frames, and similarly acquired images of continuous volumes at high contrast (supplementary text 5, fig. S5). Compared to Bessel foci, the pairs of Bessel droplets probed larger axial lengths (Fig. 2g) with much less side-ring contamination.

We also discovered that Bessel-droplet foci are more resistant to optical aberrations than Bessel foci (supplementary text 6, fig. S6). Previous investigation by us indicated that Bessel foci are degraded by Zernike aberration modes 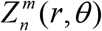 with |m| > 1 (e.g., astigmatism and trefoil) (24). In contrast, astigmatism and trefoil degraded Bessel-droplet PSFs much less than Bessel PSFs in simulation. Importantly, whereas these aberration modes strongly decreased central-peak signal and increased side-ring contribution in Bessel PSFs, their impacts on Bessel-droplet PSFs were limited to a more moderate reduction of central-peak signal without enhancing side-ring contamination. Experimentally, aberrations from our optical system degraded the peak brightness of the Bessel PSFs by 1.8–2.8× for NAs ranging from 0.4 to 0.7 and increased the relative contribution of side rings. However, the same aberration only degraded the peak signal of the Bessel-droplet PSFs by 1.03–1.08×, without any perceivable enhancement of side-ring contamination. Throughout the manuscript, system-induced aberrations were corrected for all experiments, using a novel adaptive optics procedure (24), to ensure an accurate comparison between Bessel-droplet foci and conventional Bessel foci.

### Bessel-droplet foci enable high-resolution high-contrast volumetric imaging of synapses in brain slices and in vivo

Our simulation and experimental investigations suggested that Bessel-droplet foci are uniquely suited for high-resolution and high-contrast volumetric 2PFM imaging with enhanced resistance to optical aberrations. We first tested their performance in biological tissues by imaging dendrites and dendritic spines in Thy1-GFP line M mouse neocortical slices (Materials and Methods). Using a Gaussian focus of 1.05 NA, we acquired an image stack of a 60-μm-thick volume (Fig. 3a), then imaged the same volume with 2D scanning of Bessel foci and Bessel-droplet foci of NA 0.4 and 0.7, respectively (Figs. 3b,c).

**Fig. 3.**
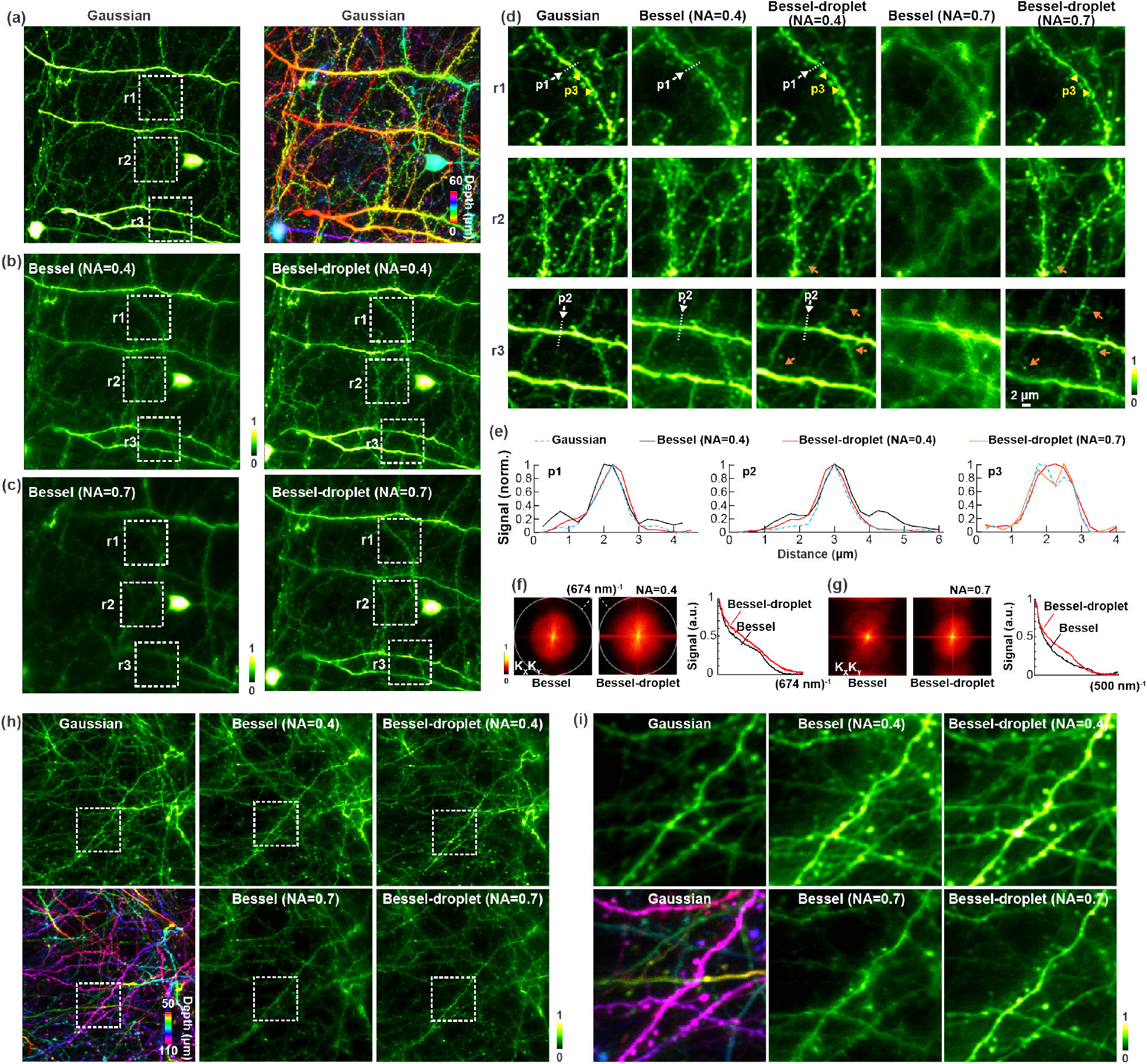
Bessel-droplet foci enable high-contrast, high-resolution volumetric imaging of synapses in mouse brain slices. (**a**) A 128 μm×128 μm×60 μm image stack of a Thy1-GFP line M brain slice, displayed in mean intensity projection and color-coded depth projection, acquired with a Gaussian focus of 1.05 NA. (**b, c**) Images acquired with Bessel and Bessel-droplet foci of 0.4 NA (**b**) and 0.7 NA (**c**). (**d**) Zoomed-in views of the regions r1, r2, r3 in **a**, **b** and **c**. (**e**) Lateral profiles of dendrites (p1 and p2) and two neighboring spines (p3) indicated by white arrows and yellow arrowheads in **d**. (**f, g**) Spatial frequency space (K_x_K_y_) representations of the images in **b** and **c**, and their radially averaged profiles. (**h**) *In vivo* imaging of neurites in the primary visual cortex of a Thy1-GFP line M mouse with Gaussian focus of 1.05 NA, Bessel and Bessel-droplet foci of 0.4 NA and 0.7 NA, respectively. (**i**) Zoom-in views of the white dashed squares in **h**. Remote focusing with an additional SLM was used to axially scan the Bessel-droplet foci in brain slice measurement; Fast axial scanning by alternating between the two phase profiles in **Fig. 2f** was used for in vivo experiment. Post-objective powers for brain slice: 22 mW for Gaussian focus, 60 mW for Bessel and Bessel-droplet foci of NA 0.4, 48 mW for Bessel and Bessel-droplet foci of NA 0.7. Post-objective powers for in vivo imaging: 88 mW for Gaussian, 100 mW for Bessel and Bessel-droplet foci of NA 0.4, 80 mW for Bessel and Bessel-droplet foci of NA 0.7. Pixel size for all images: 0.25 μm.

At 0.4 NA, both Bessel and Bessel-droplet foci allowed us to resolve and detect the dendritic spines observed in the Gaussian stack (Figs. 3b,d). However, the Bessel-droplet focus generated images with better contrast. Ghost images of dendrites, visible in the Bessel images due to side-ring excitation, were completely suppressed in the Bessel-droplet images (e.g., p1 and p2: white arrows, Figs. 3d; line profiles, Fig. 3e). At 0.7 NA, the Bessel focus gave rise to images with exceedingly poor quality, in which neither dendrites nor dendritic spines could be clearly resolved. In contrast, even at high NA, the Bessel-droplet focus generated images of similar contrast to the 0.4-NA case due to its strong side-ring suppression. The background suppression for both 0.4- and 0.7-NA Bessel droplet foci led to similar image contrast to that obtained by the 1.05-NA Gaussian focus (Fig. 3e).

The high NA also led to higher lateral resolution. The 0.7-NA Bessel-droplet focus was able to resolve neighboring spine heads (p3: between yellow arrowheads, Fig. 3d; line profiles, Fig. 3e) as effectively as the 1.05-NA Gaussian focus. It also generated much sharper images of synapses (e.g., orange arrows, Fig. 3d) than 0.4-NA Bessel-droplet focus. This resolution increase can be easily appreciated from the Fourier transforms of the full field-of-view images and their radially averaged profiles (Figs. 3f,g), with the images taken by the 0.7-NA Bessel-droplet focus having larger amplitudes for high spatial frequency components.

In addition to measurements in brain slices, we used Bessel-droplet foci of NA 0.4 and 0.7 to image volumes of dendrites and dendritic spines within a cortical depth of 50 μm to 110 μm below pia in a Thy1-GFP line M mouse brain in vivo (Fig. 3h,i). Similar to the brain slice measurements, we found that Bessel-droplet foci outperformed Bessel foci of the same NA, generating images with less side-ring excitation and higher contrast, with the highest lateral resolution obtained with Bessel-droplet focus of NA 0.7.

### Bessel-droplet foci enable *in vivo* volumetric imaging of cerebral vasculature and lymphatic circulation

The high contrast and high resolution of 2PFM using Bessel-droplet foci enabled us to resolve and monitor lymphatic vessels in the brain, which was discovered only recently to closely line the blood vessels and provide a route for surveying immune cells to circulate (25). Labeling the blood serum with dextran-conjugated Texas Red (5%, 70 kDa), we first imaged a 60-μm thick volume of the vasculature in the mouse cortex by scanning a 1.05-NA Gaussian focus in 3D (Fig. 4a,b). Whereas a single Gaussian 2D image only sampled a thin cross section of the vasculature network (inset, Fig. 4a), scanning an axially extended focus in 2D enabled rapid volumetric investigation of the same volume (Fig. 4c,d).

**Fig. 4.**
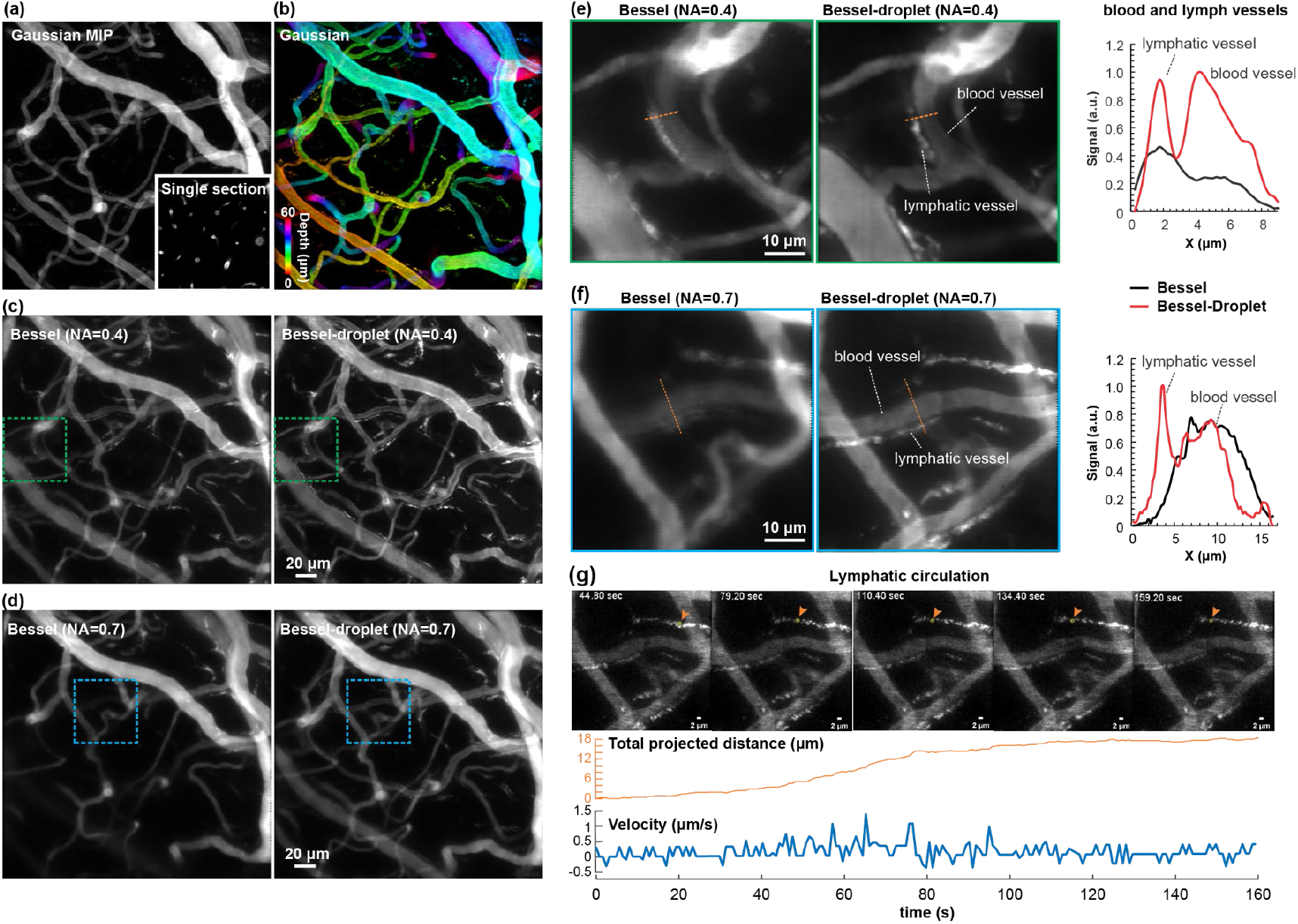
In vivo volumetric imaging of cerebral blood vessels and lymphatic circulation with Bessel-droplet 2PFM. (**a, b**) A 256 μm×256 μm×60 μm Gaussian image stack of vasculature in the mouse brain, displayed in mean intensity projection (**a**) and color-coded depth projection (**b**). Inset: a single 2D image from the stack. Pixel size: 0.5 μm. (**c, d**) Images acquired with Bessel and Bessel-droplet foci of 0.4 NA (**c**) and 0.7 NA (**d**). Pixel size: 0.5 μm. (**e,f**) (Left) Images of the volumes within the dashed boxes in c and d acquired with Bessel and Bessel-droplet foci of 0.4 NA (**e**) and 0.7 NA (**f**). (Right) Line profiles along the dashed orange lines. Pixel size: 0.25 μm. (**g**) Example frames from a 160-second recording acquired with a 0.7-NA Bessel-droplet focus. Motion of a fluorescent structure (orange arrowheads and yellow circles) within the lymphatic vessels was tracked to obtain its projected traveling distance and velocity versus time. Remote focusing with an additional SLM was used to axially scan the Bessel-droplet focus. Post-objective powers: 11 mW for Gaussian focus, 50 mW for Bessel and Bessel-droplet foci of NA 0.4, 40 mW for Bessel and Bessel-droplet foci of NA 0.7.

Both here and in our previous investigation (16), scanning the 0.4-NA Bessel focus in 2D captured all the vasculature within this volume down to capillary resolution. The 0.4-NA Bessel-droplet focus, however, provide images with better contrast and resolution (Fig. 4e), allowing the lymphatic vessels (visualized by fluorescent structures within – likely immune cells that gobbled up fluorescent dyes) to be more easily resolved from the blood vessels (Fig. 4g).

Similar to the results in the brain slice, the 0.7-NA Bessel focus generated images of the poorest quality, whereas the 0.7-NA Bessel-droplet focus gave rise to the sharpest image, in which fluorescent structures within lymphatic vessels can be easily resolved from one another (Fig. 4f). Capturing the dynamics of the lymphatic and blood vessels within this volume at 1.25 volumes per second using the 0.7-NA Bessel-droplet focus, we were able to track the movement of the fluorescent structures (supplementary movie S1). During a 160-second imaging duration, a fluorescent structure (orange arrowheads and white circles, Fig. 4g) moved a total projected distance of 17.5 μm, with intermittent backward and forward movements at velocities between −0.4 μm/s to 1.3 μm/s.

### Bessel-droplet foci enable high-sensitivity, high-contrast volumetric recording of calcium activity at synapses *in vivo*

Volumetric imaging of synaptic activity with high resolution and signal-to-noise ratio (SNR) is essential in understanding the input-output relationships in neuronal computation.

With Bessel-droplet foci, we can now measure neuronal activity at synapses at high NA. As an example, we measured the visually evoked calcium activity in synapses labeled by the calcium indicator GCaMP6s (26) in the awake mouse primary visual cortex. Presenting drifting grating stimuli of 12 different orientations (20 trials for each stimulus, Materials and Methods), we imaged synapses (i.e., dendritic spines and axonal boutons) within a 60-μm thick volume (130 – 190 μm below dura, Fig. 5a) in cortical layer II/III using Bessel and Bessel-droplet foci of 0.4 NA (Fig. 5b, supplementary movie S2) and 0.7 NA (Fig. 5c, supplementary movie S3), respectively, at 1.6 volumes/s.

**Fig. 5.**
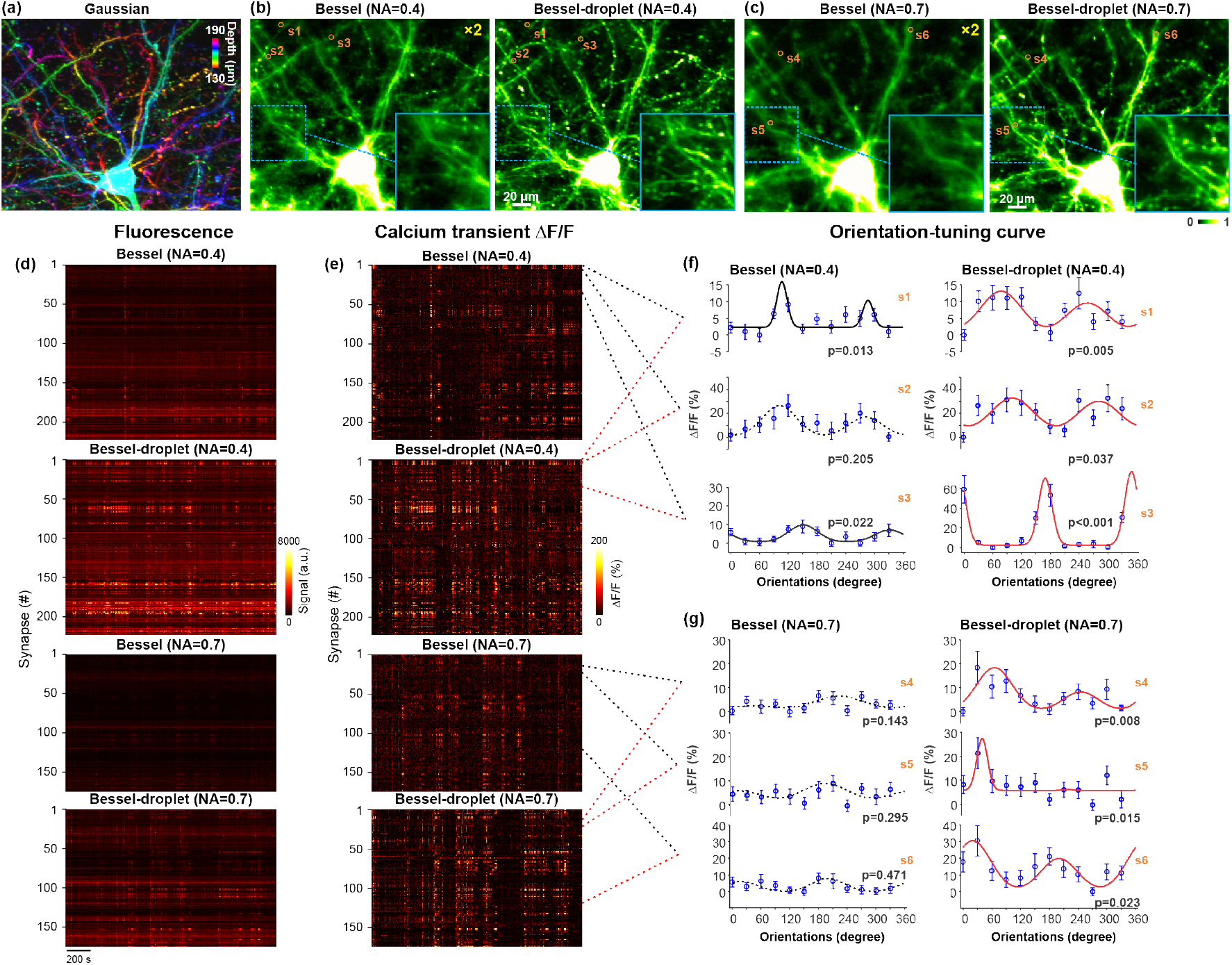
Imaging visually evoked calcium signaling at synapses with high contrast and high sensitivity in the mouse primary visual cortex in vivo. (**a**) A 128 μm×100 μm×60 μm image stack of synapses expressing GCaMP6s of a volume, displayed in color-coded depth projection, acquired with a Gaussian focus of 1.05 NA. Pixel size: 0.5 μm. (**b,c**) Images acquired with Bessel and Bessel-droplet foci of 0.4 NA (**b**) and 0.7 NA (**c**). (**d**) Raster plots of fluorescence signal traces of synapses during drifting-grating visual stimuli measured with Bessel and Bessel-droplet foci of NA 0.4 and 0.7. (**e**) Raster plots of calcium transient amplitude (ΔF/F) traces of the synapses in **d**. (**f,g**) Tuning curves for example synapses (s1-s6 labeled in **b,c**, left ends of dashed lines between **e** and **f**), obtained by fitting the averaged ΔF/F evoked by each grating stimulus with a double Gaussian function. Solid and dashed lines: tuning curves passing and not passing the statistical test (ANOVA, p<0.05) for orientation selectivity, respectively. Remote focusing with an additional SLM was used to axially scan the Bessel-droplet focus. Images taken with Bessel foci in **b** and **c** are digitally enhanced by 2× for visibility. Post-objective powers: 110 mW for Gaussian, 100 mW for Bessel and Bessel-droplet of NA 0.4, 80 mW for Bessel and Bessel-droplet of NA 0.7.

Similar to the morphological measurements of synapses, Bessel-droplet foci generated images of synapses with better contrast, and, at the same post objective excitation power, brighter fluorescence signal than Bessel foci of the same NA. They allowed us to clearly resolve synapses and accurately measure the fluorescence signal traces during visual stimulation.

Identifying the same synapses from both Bessel and Bessel-droplet datasets and comparing their time-dependent fluorescence signal (Fig. 5d), we found that Bessel-droplet excitation increased the time-averaged fluorescence of these synapses by 1.68±0.55 fold for NA 0.4 (mean±S.D., N=223 synapses) and 1.84±0.32 fold (N=175 synapses) for NA 0.7, respectively. Calculating the calcium transient amplitude as the percentage of fluorescence signal change (ΔF/F) (Fig. 5e), we found that Bessel-droplet foci increased the time-averaged calcium transient amplitude by 1.30±0.34 fold for NA 0.4 and 1.52±0.36 fold for NA 0.7, respectively, due to less contamination of unresponsive neuropils through side-ring excitation.

This increase in signal and contrast led to an increase in SNR, which enabled us to detect orientation selectivity from more synapses at both NA 0.4 (Fig. 5f) and NA 0.7 (Fig. 5g). Whereas the tuning curves measured by Bessel foci for many synapses failed to pass the statistical test used to detect orientation selectivity (dashed tuning curves, Fig. 5f and 5g), their tuning curves obtained with Bessel-droplet foci (solid tuning curves, Fig. 5f and 5g) passed the test (ANOVA, p<0.05 for orientation-selective synapses, Materials and Methods). Out of 223 synapses identified from the NA-0.4 datasets, Bessel and Bessel-droplet foci detected 5 and 25 orientation-selective synapses, respectively. Out of 175 synapses identified from the NA-0.7 datasets, Bessel and Bessel-droplet foci detected 2 and 9 orientation-selective synapses, respectively. (Fewer synapses were observed for NA 0.7 due to their shorter axial extension, as indicated by Fig. 2g.)

## Discussion

Here, we described a method to suppress the side-ring excitation in 2PFM when using axially extended focus such as a Bessel beam. Given that an annular illumination pattern generates a Bessel-like focus, we shaped the excitation light into two concentric annuli at the objective back pupil plane, generating two Bessel foci that interfered to suppress energy distributed in the side rings and formed droplet-like excitation pattern axially.

We developed a computational approach that led to phase profiles that, when applied to the wavefront of the excitation laser using a SLM, allowed us to generate Bessel-droplet foci at high efficiency. Previously in single-photon fluorescence light-sheet microscopy (27), an amplitude mask was used to transmit two concentric annuli to generate Bessel-droplet foci with suppressed side-ring fluorescence excitation for high-contrast imaging. Besides having a much lower power throughput than our SLM-based method, the amplitude mask alone cannot shift droplet locations along its propagation direction. As a result, only the area illuminated by a single droplet was imaged, severely limiting the field of view size. Using wavefront engineering through the SLM, we were also able to control the relative phase between the two illumination annuli, which shifted the axial locations of the droplets to sample volume continuously. Our approach thus allowed us to utilize the excitation by multiple droplets to probe a much larger volume. Together with its high throughput, our method is also ideally suited for single-photon fluorescence microscopy modalities including in the fluorescence emission path for high-contrast extended depth-of-field widefield imaging (28).

Generating Bessel-droplet foci from 0.4 to 0.7 NA, experimentally we observed up to 10.6 fold suppression of side-ring two-photon fluorescence compared to Bessel foci of the same NAs. Higher NA Bessel-droplet can also be generated, with the upper limit on NA posed by SLM, which need to have sufficient pixel resolution to display the phase profile at high fidelity. Translating the droplet excitation axially either using a defocus pattern remotely applied by another SLM or, more simply, switching between phase profiles leading to 0 or π phase offset of the two illumination annuli, we can continuously probe volumes with even larger axial extents than their Bessel counterparts. With much stronger side-ring suppression, much higher excitation efficiency, and much longer axial extent, our method is superior to an alternative approach that relied on incoherent addition of multiple Bessel beams to smooth out (but not suppress) side-ring excitation (29). Compared with a recent study that removed side-ring contamination by subtracting the image taken with a 3rd-order Bessel beam from the image taken with a 0th-order Bessel beam (30), our method is more efficient in excitation and causes less photobleaching by suppressing side-ring excitation altogether rather than removing its contribution via post processing. Our method can also be used to study more weakly fluorescent samples, whose images have low SNR and would make the subtraction-based approach infeasible.

We validated the superior performance of Bessel-droplet foci by using them for extended depth-of-field imaging in both brain tissue and the awake mouse brain in vivo. At both 0.4 and 0.7 NA, Bessel-droplet foci generated images of synapses, capillaries, and lymphatic vessels with higher contrast. For subcellular, synapse-resolving imaging in the brain at 0.7 NA, the Bessel-droplet focus represented the only viable option, with the side-ring excitation from the 0.7-NA Bessel focus yielding images of much poorer quality. Compared to 0.4-NA Bessel foci used in previous studies, 0.7-NA Bessel-droplet foci improved the lateral spatial resolution by 1.6×, allowing us to resolve neighboring spines. The higher spatial resolution and superior side-ring suppression of Bessel-droplet foci also proved beneficial for functional imaging, giving rise to better sensitivity in capturing calcium transients of larger amplitudes and a better accuracy in orientation tuning characterization. Enabling high-NA operation, Bessel-droplet foci represent an attractive approach for high-contrast, high-resolution volumetric 2PFM imaging in biological tissue.

## Materials and Methods

### Animal preparations

All animal experiments were conducted according to the National Institutes of Health guidelines for animal research. Procedures and protocols on mice were approved by the Institutional Animal Care and Use Committee at the University of California, Berkeley. Brain slices of 200 μm in thickness from a Thy1-GFP mouse line M (male, > 2months old) were used for dendrites and dendritic spine imaging. A wild-type mouse (C57BL/6J, male, > 2months old) was used for lymphatic circulation imaging. The mouse was head-fixed under the microscope objective and anesthetized with isoflurane. 50 μL of 5% (w/v) 70-kDa dextran-conjugated Texas Red fluorescence dye solution was then injected into the animal’s retro-orbital sinus. For visual stimulation evoked calcium signaling imaging, we used a wild-type mouse (C57BL/6J, male, > 2months old) with sparse expression of GCaMP6s in the primary visual cortex at layer II/III. The sparse expression was achieved by injection of 20 nL 1:1 mixture of diluted AAV2/1-syn-Cre virus (original titer: 1012 infectious units per mL, diluted 1,000-fold in phosphate-buffered saline) and Cre-dependent GCaMP6s (AAV2/1.syn.Flex.GCaMP6s, 8 × 1011 infectious units per ml) at a depth between 150 μm to 200 μm below the pial. Cranial window implantation for both vasculature imaging and calcium activity imaging was carried out following procedures described previously (13, 31).

### Visual stimulation and data analysis

Visual stimulation was delivered by a blue LED light source (450–495 nm, SugarCUBE) that was back-projected on a screen (Teflon film, McMaster-Carr) using a customized projector (32). The screen was positioned 17 cm from the right eye, covering 75°× 75° of visual space and oriented at ~40° to the long axis of the animal. The visual stimulation was presented as full-field gratings drifting in 12 directions (0° to 330° at 30° increments) in pseudorandom sequences. Gratings contrasts were set to be 100% with a spatial frequency of 0.07 cycles per degree and a temporal frequency of 26° s-1. Each grating stimulus lasted 6 sec interleaved with 6 sec gray screen (4 sec before and 2 sec after the grating stimulus). Each stimulus was repeated for twenty trials.

We identify regions of interest that likely to be synaptic structures (e.g., dendritic spines or axonal boutons) and extract the averaged fluorescence brightness of their pixels for each frame to obtain signal trace for each synaptic structure during visual stimulation. We defined the basal fluorescence *F_0_* to be the averaged signal across 4 sec of blank period prior to grating presentation. The amplitude of the calcium transients *R* was calculated as the averaged *ΔF*/*F_0_* across the 6 sec of grating stimulus. Drifting-grating-evoked calcium responses were defined as orientation-selective if the calcium transient amplitudes for the 12 grating stimuli were significantly different (one-way ANOVA, p < 0.05). Assuming a normal distribution of *R* and equal variance across grating direction *θ*, the response evoked by drifting gratings of direction *θ* can be fitted with a bimodal Gaussian function (33):

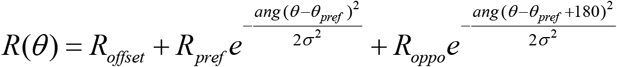

where *R_offset_* is a constant offset, *R_pref_* and *R_oppo_* are the responses at the preferred grating drifting angle *θ_pref_* and *θ_pref_* - 180°, respectively.

## Acknowledgements

We thank L. Abdeladim for an inspiring discussion and S. Temprana of the Adesnik laboratory for providing test samples.

## Funding

National Institutes of Health grant U01NS103489 (NJ)

National Institutes of Health grant 2R44MH111463 (NJ)

National Institutes of Health grant RF1MH120680 (HA)

## Author contributions

NJ supervised the project. NJ and WC conceptualized the project. WC developed the computational method and built the Bessel-droplet module. QZ, RN, and FJ prepared samples. WC collected and analyzed data. WC and NJ wrote the manuscript. All authors reviewed the manuscript.

## Competing interests

The authors declare that there are no conflicts of interest related to this paper.

## Data and materials availability

All data are available in the main text or the supplementary materials. Additional data related to this paper may be requested from the authors.

## Notes

### Competing Interest Statement

The authors have declared no competing interest.

