## Supplemental materials for "Bessel-droplet foci enable high-resolution and high-contrast volumetric imaging of synapses and circulation in the brain *in vivo*"

### This PDF file includes:

Supplementary Methods and Analyses 1-6
Fig. S1. Computing phase patterns for generating two concentric annular illumination.
Fig. S2. Optical throughput comparison between the SLM-based approach and the amplitude-mask-only approach for Bessel-droplet foci of different numerical apertures.
Fig S3. Optimizing concentric annular illumination for maximal side-ring suppression of Bessel-droplet foci.
Fig.S4. Measured axial PSFs of Bessel droplets of NA 0.4, 0.5, 0.6, 0.7 without and with fast axial scanning (FAS).
Fig S5. Axial scanning of Bessel droplets with remote focusing by a SLM conjugated to the objective pupil.
Fig S6. Bessel-droplet foci are more resistant to optical aberrations than Bessel foci.

**1. Computing phase patterns for generating two concentric annular illumination.**

We developed a procedure of computing the phase patterns on the SLM (see Fig. 1a of the main text for optical setup) to generate two concentric annuli with specified phase offset $\Delta\phi$ at the mask/objective back pupil plane. In Step 1 (Fig. S1a), we started with the desired optical field distributions at the objective back pupil plane, for example, two concentric annuli (radii: $r_1$ and $r_2$; thickness: $d$; Fig. S1a1) with phase differences ($\Delta\phi$) between the two annuli being either 0 (Fig. S1a2) or $\pi$ (Fig. S1a3). In Step 2, we inverse Fourier transformed the field distribution using customized MATLAB® codes (Supplemental Code). The result was a complex optical field (Fig. S1b), with its phase pattern (e.g., Fig. S1b2 and S1b4) displayed by the phase-only SLM for Bessel droplet generation. In Step 3, we validated the above method by Fourier transforming the phase pattern to obtain the resulting electric field distribution at the mask/pupil plane, which indeed had the desired two-annular amplitude profile and correct phase offset (Fig. S1c).

**2. Optical throughput comparison between the SLM-based wavefront shaping approach and the amplitude-mask-only approach.**

An amplitude mask with two concentric transmissive annuli, as used in our system, can be directly placed in the Gaussian beam path to generate concentric annular illumination, as previously used in single-photon fluorescence imaging modalities (e.g., wide field microscopy (28) and light-sheet microscopy (27). Here we compared experimentally the optical throughput of our SLM-based wavefront shaping approach with this amplitude-mask-only approach for Bessel-droplet foci of different NA. For our SLM-based module, 38%, 36%, 36%, and 35% of the power incident on the SLM was detected after the mask for Bessel-droplet foci of NA 0.4, 0.5, 0.6, and 0.7, respectively. In comparison, the amplitude-mask-only approach had optical throughputs of 6%, 7%, 8% and 8%, respectively, substantially lower than the SLM-based approach.

**3. Optimizing concentric annular illumination to maximize side-ring suppression for Bessel-droplet foci.**

For a conventional Bessel focus, larger fraction of focal energy is distributed to the side rings with higher NA. We simulated this effect by calculating two-photon excitation PSF using a vectorial diffraction model developed by Richards & Wolf (23). We first characterized the ratio between the peak of the most prominent side ring and the central peak of the lateral PSF, which showed an increase with NA for Bessel foci (Fig. S3a). For Bessel-droplet foci, we fixed the radius of the outer annulus ($r_1$) and annular thickness ($d$) but varied the inner annulus radius ($r_2$). When $r_2/r_1$=1, we obtained the side-ring ratios for Bessel foci. The side-ring ratios of Bessel-droplet foci of various NAs exhibited multiple local minima with the increase of $r_2/r_1$ (Fig. S3b), with the global minima appearing at $r_2$/r$_1$ between 0.5 to 0.6 for different NA (indicated by the dashed vertical lines), which were used as the optimal designs for the concentric annulus for Bessel-droplet foci. This side-ring contamination was further quantified as the ratio between the integrated signal within the most prominent side ring and the central peak of the lateral PSF, which as expected showed an increase with NA for Bessel foci (Fig. S3c), and the greatest side-ring suppression at $r_2$/r$_1$ ~ 0.6 for Bessel-droplet foci (Fig. S3d). As expected, the Bessel-droplet foci generated by concentric annular illuminations with the optimum $r_2/r_1$ showed strong suppression of side rings in their simulated lateral and axial PSFs (Fig. S3e).

**4. Fast axial scanning of Bessel-droplet foci by alternating the phase profiles on the high-speed SLM.**

For fast axial scanning of Bessel droplets, the high-speed SLM alternated between the two phase profiles that generated two concentric annular illumination with the phase offset being

0 or π for consecutive frames. This created an axially continuous sampling of volumes (Fig. S4).

**5. Axial scanning of Bessel-droplet foci with remote focusing by a SLM conjugated to the objective back pupil plane.**

In addition to the fast focus scanning described above, using the additional SLM in our 2PFM (one that is conjugated to the objective back pupil plane, SLM2, Fig. S5), we can axially scan the Bessel-droplet foci by applying a defocus pattern on SLM2. We used both methods in taking data from biological samples.

**6. Bessel-droplet foci are more resistant to optical aberrations than Bessel foci.**

We used a numerical model based on the vectorial diffraction theory derived by Novotny and Hecht (34) to calculate PSFs for pupil electric fields with noncircularly symmetric wavefront aberrations (e.g., astigmatism and trefoil). Applying the wavefront distortion of Zernike modes corresponding to astigmatism ($Z_2^2$), trefoil ($Z_3^3$), and coma ($Z_3^1$) of peak-to-valley value of 1 wavelength at the pupil plane, we simulated the PSFs of Bessel and Bessel-droplet foci. As shown in the lateral PSFs and their line cross sections (Fig. S6a,b), astigmatism and trefoil severely decreased the peak signals and enhanced the side-lobe contribution of the Bessel focus, but only moderately decreased the peak signal of the Bessel-droplet focus without increasing the side-ring contribution.

Experimentally, we compared the impact of optical system aberrations on Bessel and Bessel-droplet foci of NA=0.4, 0.5, 0.6 and 0.7. Using a highly efficient and effective aberration correction method, where corrective wavefront measured at the objective pupil plane was computationally propagated to the focal plane SLM (24) so that we can correct for both amplitude and phase distortion at the objective pupil plane (Fig. S6c), we measured the PSFs without and with system aberration correction (Fig. S6d,e). For Bessel foci, system aberration decreased peak brightness of the PSF and increased the relative contribution of side-ring contamination. In contrast, system aberration only reduced peak brightness minimally without any perceivable enhancement of side-ring contamination.

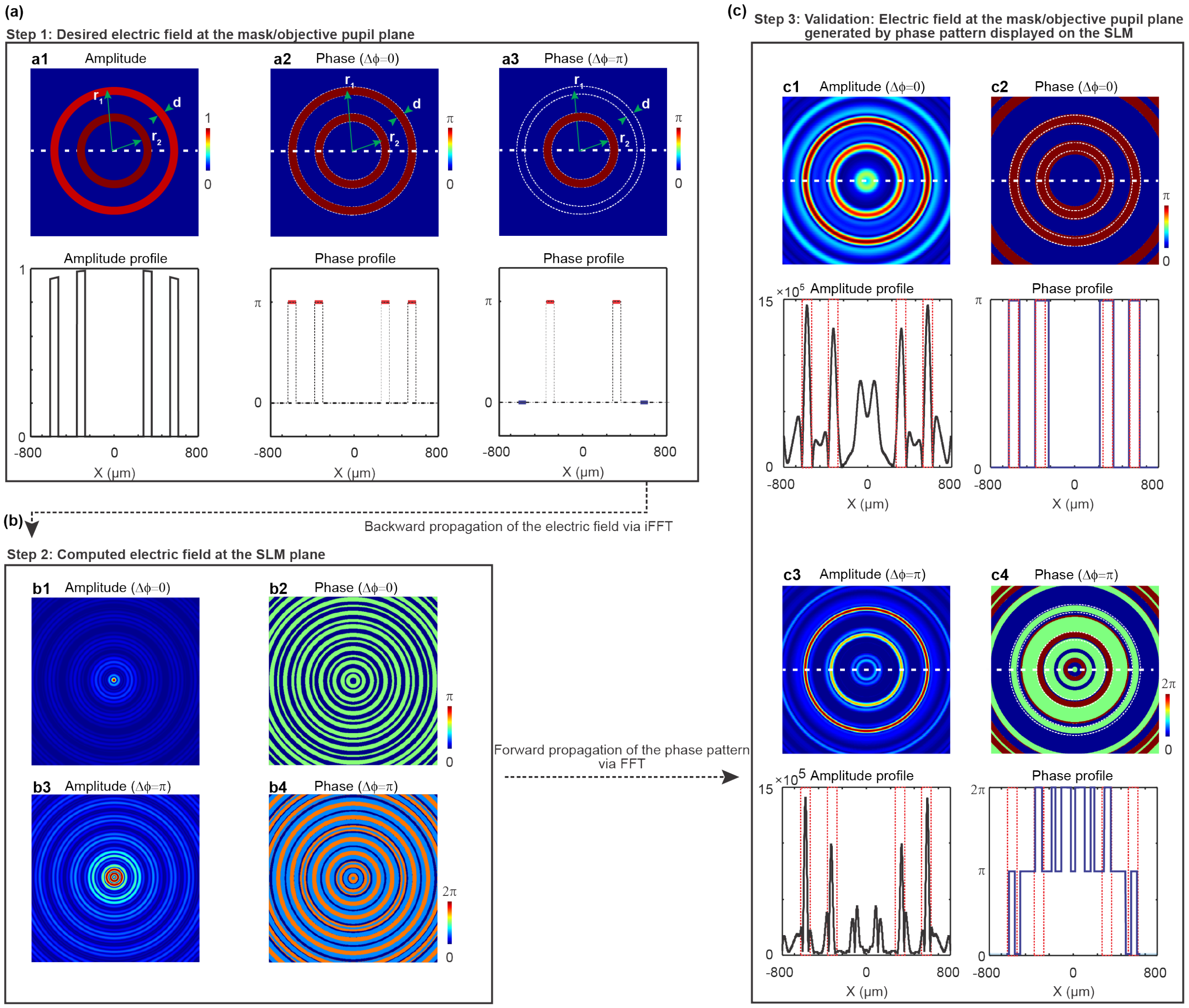


**Fig. S1. Computing phase patterns for generating two concentric annular illumination.** (**a**) Desired electric fields at the mask plane. Electric field at the objective back pupil plane is scaled by the magnification factor 4.667. **a1:** Amplitude distribution of the two annuli. **a2**, **a3:** Phase patterns with the phase offset between the two annuli $(\Delta\phi)$ being 0 or $\pi$. (**b**) Inverse Fourier transforming the electric field distributions in **a** to obtain the amplitude (**b1**,**b3**) and phase (**b2**,**b4**) patterns of the electric field at the SLM plane. Experimentally, the phase patterns (**b2**,**b4**) are displayed on the phase-only SLM. (**c**) Validation: Amplitude (**c1**,**c3**) and phase (**c2**,**c4**) patterns of the electric field at the mask/objective pupil plane, generated by displaying **b2** and **b4** on the SLM, respectively. Vertical red dashed lines indicate the transmissive areas of annular mask. Design dimensions on the mask: $r_1$=0.617 mm, $r_2$=0.311 mm, $d$=64 μm, NA=0.4, Axial FWHM of Bessel-droplet is 44 μm. MATLAB codes for Step 1-3 are provided in **Supplementary Codes**.

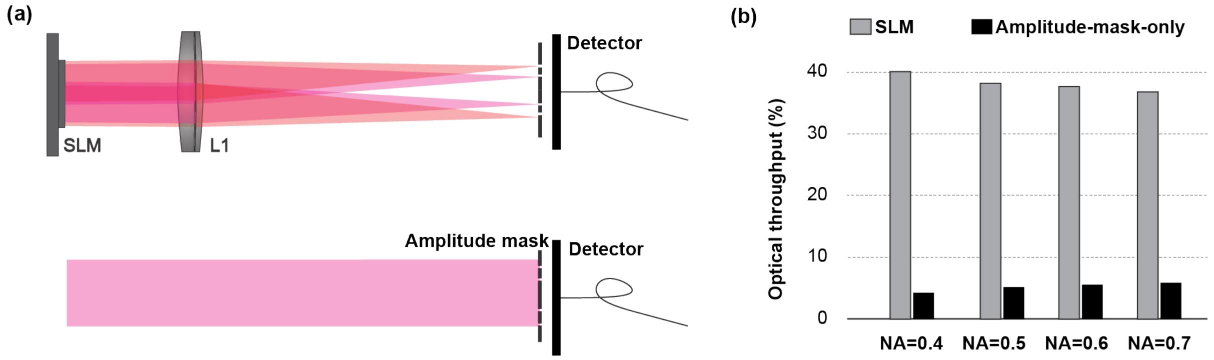


**Fig. S2. Optical throughput comparison between the SLM-based approach and the amplitude-mask-only approach for Bessel-droplet foci of different numerical apertures.** (**a**) Schematics of the SLM-based module and the amplitude-mask-only module. (**b**) Optical throughputs of the two approaches for Bessel-droplet foci of NA 0.4, 0.5, 0.6, and 0.7.

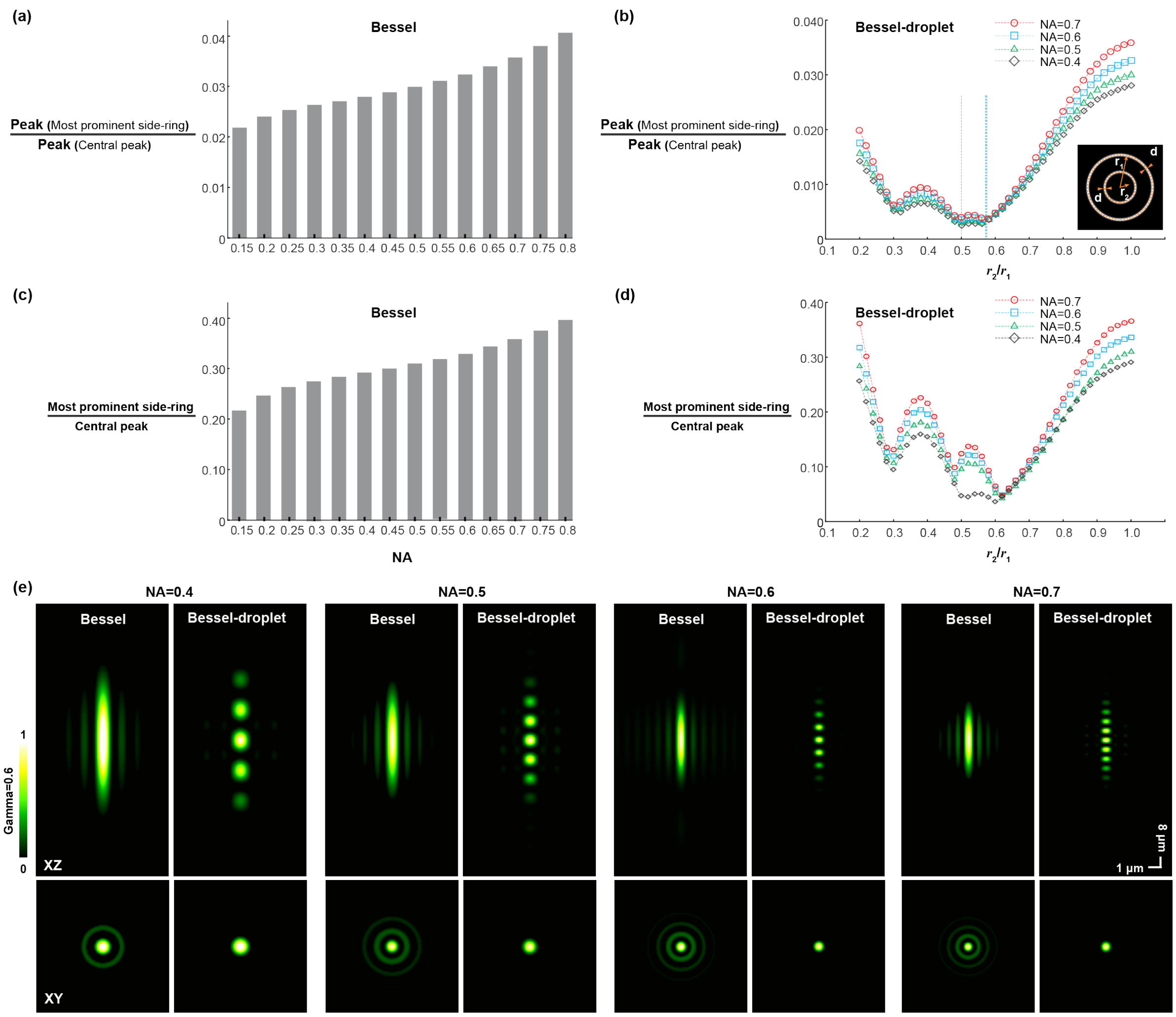


**Fig S3. Optimizing concentric annular illumination for maximal side-ring suppression of Bessel-droplet foci.** (**a**) Ratios between the peak of the 1st side ring and the central peak of the lateral two-photon excitation PSF for Bessel foci of different NAs. (**b**) Ratios between the peak of the most dominant side ring and the central peak of the lateral two-photon excitation PSF for Bessel-droplet foci of different NAs as a function of $r_2/r_1$. $r_1$: radius of the outer annulus; $r_2$: radius of the inner annulus. Dashed lines: locations of minimal ratio. (**c**) Side-ring contamination for Bessel foci of different NAs, calculated as the ratio between the signal integrals of the 1st side ring and the main peak of the lateral two-photon excitation PSF. (**d**) Side-ring contamination for Bessel-droplet foci of different NAs, calculated as the ratio between the signal integrals of the most dominant side ring and the main peak of the lateral two-photon excitation PSF, as a function of $r_2/r_1$. (**e**) Simulated axial and lateral two-photon excitation PSFs for Bessel versus Bessel-droplet foci of NA 0.4, 0.5 0.6, and 0.7. Gamma correction at 0.6 was applied to images to make side ring more visible.

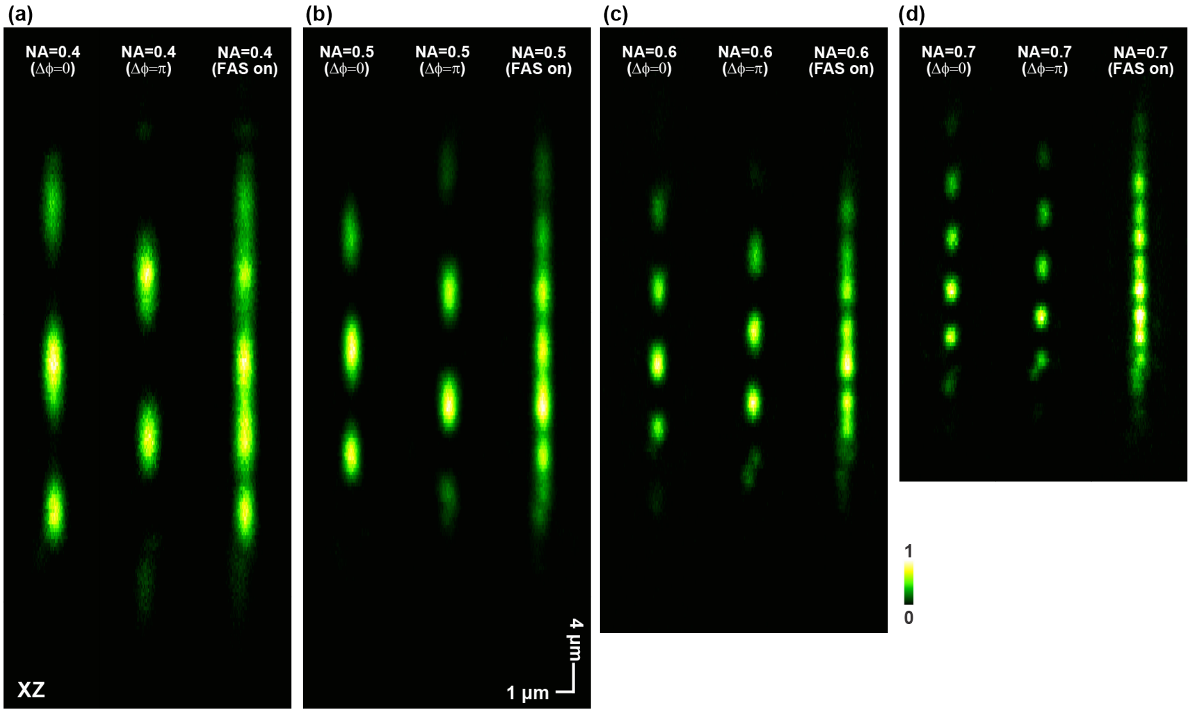


**Fig. S4. Measured axial PSFs of Bessel droplets of NA 0.4, 0.5, 0.6, 0.7 without and with fast axial scanning (FAS).** From left to right: PSF produced by a phase profile on the SLM that generates two concentric annular patterns at the back focal plane with phase offset Δϕ being 0; PSF produced by a phase profile on the SLM that generates two concentric annular patterns at the back focal plane with phase offset Δϕ being π; PSF measured while rapidly alternating the phase profile on the SLM between the previous two profiles. Measurement done on 0.2-μm-diameter beads.

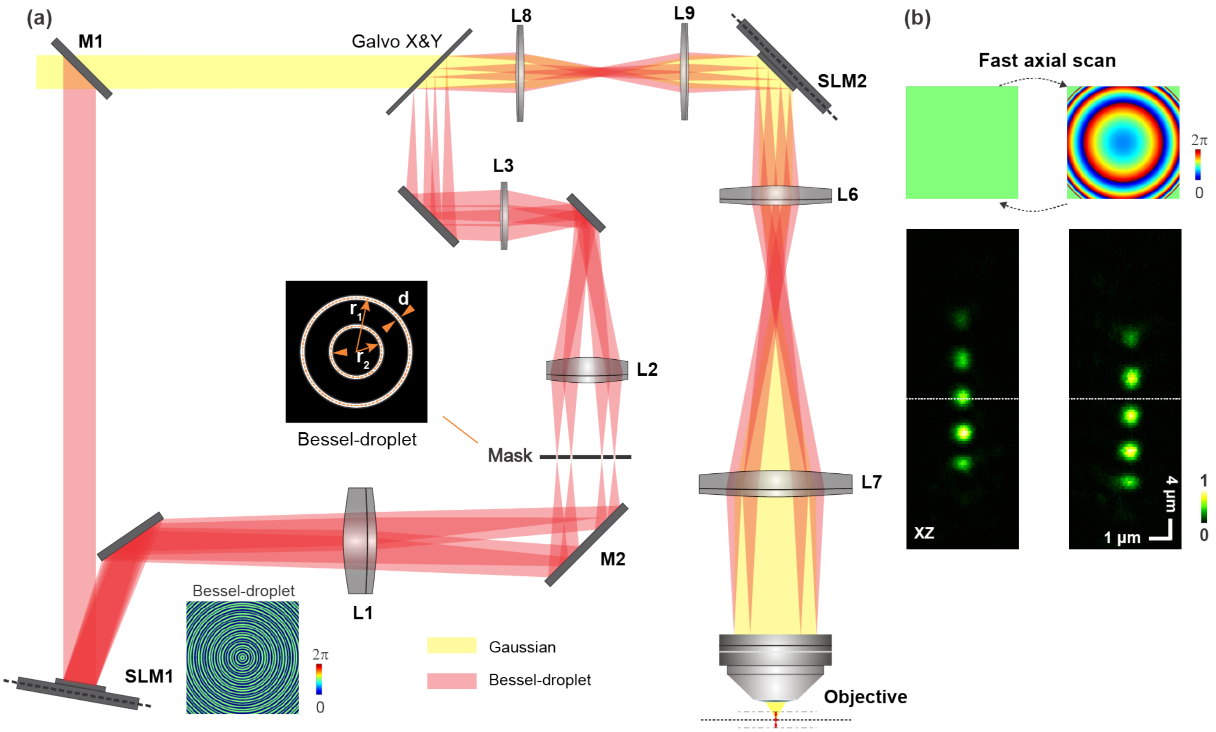


**Fig S5. Axial scanning of Bessel droplets with remote focusing by a SLM conjugated to the objective pupil.** (**a**) Optical path diagram of the 2PFM, showing the additional SLM (SLM2) that is conjugated to the objective back pupil plane. Insets: (Lower) phase pattern on the Bessel-beam-generating SLM (SLM1) and (Upper) the corresponding excitation electric field amplitude pattern at the mask/objective back pupil plane for a Bessel-droplet focus. (**b**) (Top) Two phase profiles (a flat wavefront or a defocus wavefront) displayed on SLM2 axially scans the Bessel-droplet focus as indicated by (Bottom) the axial PSFs of the Bessel-droplet foci measured with a 0.2-µm-diameter bead.

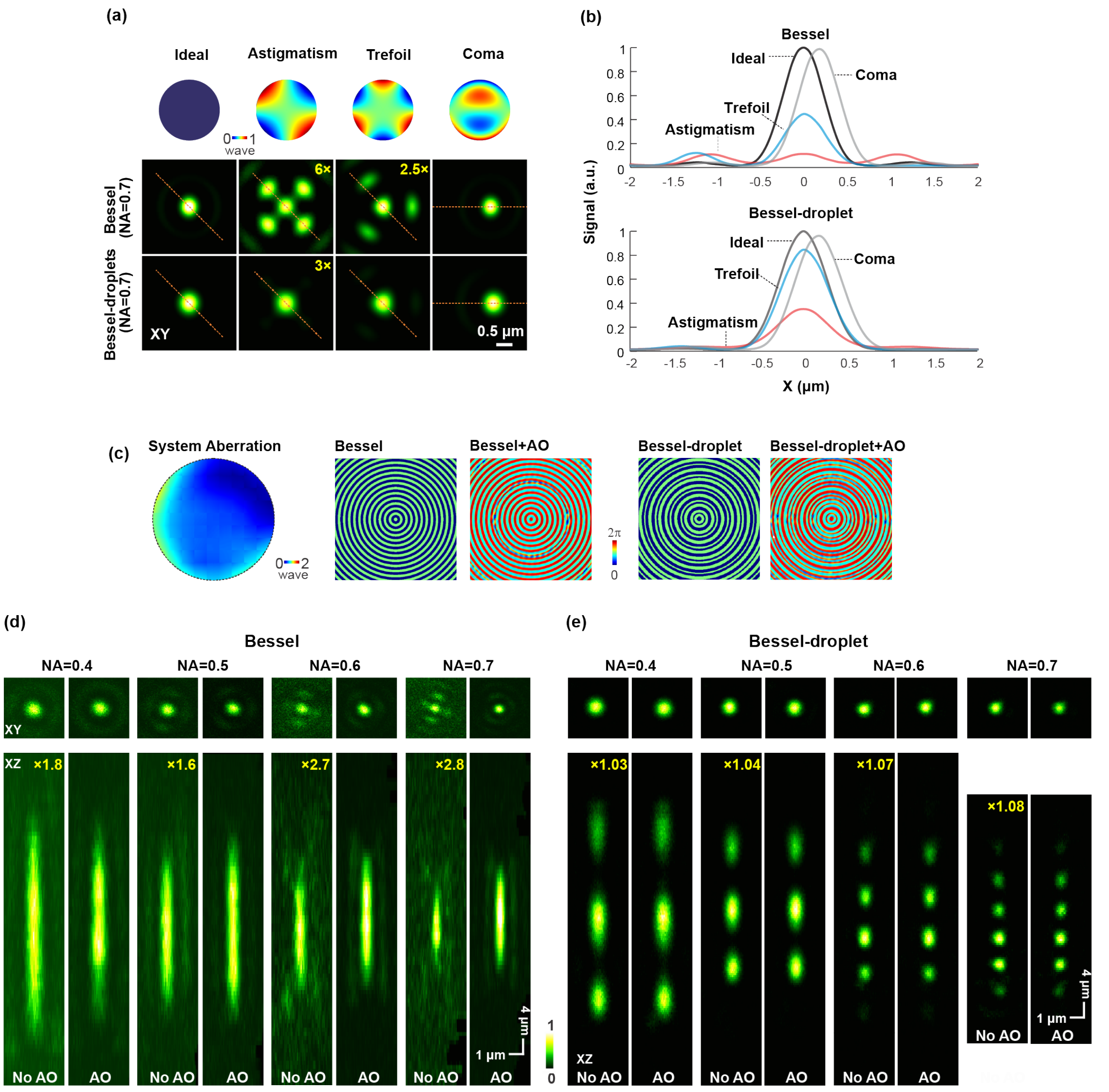


**Fig S6. Bessel-droplet foci are more resistant to optical aberrations than Bessel foci.** (**a**) Zernike modes representing astigmatism, trefoil, and coma, respectively, and their impact on the lateral (XY) PSFs of simulated Bessel and Bessel-droplet foci (NA 0.7). (**b**) Comparison of lateral PSF cross sections (along the dashed orange lines). (c) Experimentally measured system aberration, and the phase profiles on the SLM for PSFs measured without and with aberration correction. (**d**) Lateral and axial (XZ) PSFs of Bessel foci of NA 0.4, 0.5, 0.6, and 0.7 measured using a 0.2-µm-diameter fluorescent bead. (**e**) Lateral and axial PSFs of Bessel-droplet foci. In **a,d,e**, some aberrated PSFs have their signal digitally enhanced to improve visibility with the enhancement factors in orange font.
